# Causal Language Detection using Text-Document Features: Methodology and Insights from 10 Years of Gut Microbiome Research

**DOI:** 10.64898/2026.03.02.709039

**Authors:** Albina Tskhay, Cristina Longo, Alibek Moldakozhayev, Nia Kang, Celia Greenwood, Roxana Behruzi, Stan Kubow, Tibor Schuster

**Affiliations:** Department of Family Medicine, McGill University, Montréal, Québec, Canada; Centre de Recherche du CHU Sainte-Justine, Montréal, Québec, Canada; Faculté de Pharmacie, Université de Montréal, Montréal, Québec, Canada; Division of Genetics, Department of Medicine, Brigham and Women’s Hospital, Harvard Medical School, Boston, MA, USA; Department of Neurology and Neurosurgery, McGill University, Montreal, QC, Canada; Department of Human Genetics, McGill University, Montreal, Quebec; Faculty of Medicine and Health Sciences, McGill University, Montreal, QC, Canada; Outaouais Birth Center, Research Center of CISSS Outaouais, Gatineau, QC, Canada; School of Dietetics and Human Nutrition, McGill University, Montreal, Quebec, Canada

## Abstract

Detecting causal language in scientific literature is critical for understanding how research fields frame evidence and inform interventions and policies, yet existing approaches commonly rely on manual annotation. The objective of this study was to evaluate four classifiers for detecting causal language and to apply the best-performing model to assess trends in microbiome research. Microbiome research, with its rapidly expanding observational literature, provides a relevant case study. We extracted Term Frequency–Inverse Document Frequency (TF-IDF) features from the last three sentences of available publication abstracts and trained four classifiers (L1- and L2-regularized logistic regression, Random Forest, and eXtreme Gradient Boosting) to detect causal language. A total of 475 sentences, as determined pragmatically based on annotation feasibility and observed stabilization of model performance, were manually labeled as causal or non-causal following established guidelines for systematic evaluation of causal language in observational health research. Of these, 75% of sentences were used for training and 25% for testing. L1-regularized logistic regression achieved the highest performance (accuracy 76%, F1 72%, prevalence detection accuracy 95%, sensitivity 72%, and specificity 80%) and was applied to 20,022 human gut microbiome abstracts published between 2015 and 2025 grouped into 20 thematic topics using structural topic modeling. Predicted causal language prevalence declined from 52% to 44% between 2015 and 2018, then rose to 51% by 2025, with notable variation across topics (range: 43.1–53.3%). Temporal trends differed across subfields, with increases in *Metabolic disorders, Fecal microbiota transplantation, and decreases in Biomarkers and prediction, Antibiotic resistance, and In vitro fermentation*. Analysis of influential words confirmed that causal meaning is primarily driven by verbs and modifiers lexically signaling change or intervention. The proposed approach for identifying causal claims in scientific abstracts enables systematic and automated, scalable assessment of how evidence is framed. Its application to the microbiome field highlighted heterogeneity in the reporting of causal relationships and informing the interpretation of microbiome findings for clinical and public health decision-making.

## INTRODUCTION

Causal language refers to the explicit or implicit suggestion that one event or factor directly or indirectly influences another (1). Its use shapes how research questions are framed, how findings are interpreted, and how evidence is communicated. The appropriate use of causal language is especially critical in fields like medicine and public health, where implications drawn from research frequently aim at informing interventions and impact clinical decisions, policy-making, and patient outcomes (2). At the same time, causal language in the literature is often tempered by concerns over the risks of overstating what the underlying study design can support (3). There is general agreement that causal questions and hypotheses should be articulated transparently, provided that underlying assumptions, methodological limitations, and sources of uncertainty are clearly acknowledged (4, 5).

Several prior works have evaluated causal language in scientific writing using structured manual annotation. For instance, the 2018 “Causal Language and Strength of Inference in Academic and Media Articles Shared in Social Media” (CLAIMS) project examined causal language and strength of inference in social media coverage and a sample of highly cited biomedical papers, introducing a standardized review tool for assessing causal statements (2). Building on this work, a 2022 study reviewed 1,170 observational biomedical research articles to assess causal language and the strength of associated recommendations, producing a publicly available guide for manual assessment of causal claims (4). These studies provide important benchmarks for understanding how causal assertions are framed but they rely on intensive human review, limiting scalability.

In response to this limitation, several natural language processing approaches have been proposed to automate causal language detection. For instance, a linguistics-grounded framework using corpus lexicography was developed to identify causal constructions in newspaper texts, but required extensive manual annotation and validation of syntactic patterns using a hand-crafted lexicon (6). Similarly, a supervised learning model introduced in 2019 achieved high predictive accuracy for classifying causal statements, but depended on a large manually labeled training set of approximately 3000 articles (7). While these approaches demonstrate the feasibility of automated detection, many were not tailored to biomedical abstracts and require either domain-specific linguistic resources (6) or substantial annotation effort (7), constraining their applicability for large-scale literature analysis.

Microbiome research provides a particularly relevant context in which the utility of causal language detection and mapping can be examined . Since the launch of the Human Microbiome Project in 2007 (8), the field has expanded rapidly, implicating microbial communities in a wide range of health conditions, including inflammatory bowel disease, obesity, cardiovascular disease, and mental health disorders (9-11). Despite these developments, translation of microbiome findings into clinical practice remains limited, in part due to uncertainty surrounding causal interpretation: many studies rely primarily on associational analyses with limited integration of formal causal inference frameworks (12), leaving ambiguities about microbiome-host causal relationships (13, 14).

Examining how causal language is used in microbiome publications can help contextualize the field’s interpretive norms, while also serving as a test case for scalable automated detection. To this end, we propose a scalable machine learning–based approach that minimizes annotation requirements while maintaining adequate performance for corpus-level analysis. We compare four classifiers: L1- and L2-regularized logistic regression, Random Forest, and eXtreme Gradient Boosting (XGBoost), using Term Frequency–Inverse Document Frequency (TF-IDF) textual representations (15). These methods balance interpretability, robustness in high-dimensional feature spaces, and the ability to capture non-linear patterns. Regularized logistic regression provides stability and feature selection in sparse settings (16, 17), while ensemble methods such as Random Forest and XGBoost can model complex non-linear relationships (18-20).

We challenge the assumption that extensive manual labeling is necessary for reliable detection by evaluating whether standard supervised text classification models can achieve adequate performance when trained on a relatively small, domain-specific labeled dataset. Accordingly, the objectives of this study were to (a) develop and evaluate an automated approach for detecting causal language using a small, labeled dataset, and (b) apply this method to a large corpus of microbiome publications to characterize temporal and topic-specific patterns in the use of causal language.

## METHODS

### Data collection and preprocessing

We retrieved abstracts of gut microbiome-related research articles from PubMed using the easyPubMed package in R (21) . The inclusion criteria were: a date of publication between January 1, 2015, and July 23, 2025; indexing with the MeSH term “Gastrointestinal Microbiome”; human subjects; English-language publication; and abstract availability (12). Review articles, meta-analyses, case reports, editorials, and letters were excluded. Abstracts and titles were separately extracted and preprocessed by converting all text to lowercase, removing punctuation and numeric characters, and eliminating standard English stop words (22).

### Causal language classification

#### Manual annotation

To train and evaluate the causal language classification models, we manually annotated sentences extracted from the conclusion sections of microbiome abstracts. For the first 205 sentences, we used the main conclusion sentences identified in our prior methodological review of observational microbiome studies (12). Here, “main conclusion” refers to one of up to three concluding sentences that summarize the primary finding of a study. The remaining 270 sentences were randomly selected from the last three sentences of identified abstracts, ensuring coverage of diverse study conclusions, because the last three sentences were assessed to capture possible variation in how authors phrase conclusions (12). Extracted sentences were independently labeled as causal or non-causal by two reviewers following established guidelines for systematic evaluation of causal language in observational health research (4). Reviewers assessed each sentence by identifying the main exposure and outcome, considering linking words/phrases (e.g., ‘associated with,’ ‘had higher’) and modifying terms indicating direction, strength, uncertainty, or statistical interpretation (e.g., ‘may be,’ ‘significantly’). Discrepancies were resolved through discussion to reach consensus, and the resulting labels were used to train and evaluate the causal language classification models.

#### Feature construction

Sentences were transformed into numerical representations using a word-level Term Frequency–Inverse Document Frequency (TF-IDF) vectorizer. TF-IDF assigns high weights to terms that are frequent in a given sentence but rare in the overall corpus (i.e., entire text population), thereby emphasizing lexically informative patterns while down-weighting ubiquitous words (23). Sentence-level features were constructed using n-grams ranging from 2 to 5 words (24). Vocabulary pruning excluded terms with fewer than two occurrences across the corpus and those present in more than 75% of documents, reducing noise from overly rare or overly common features (25). The resulting document–term matrices served as input for model training. To prevent data leakage, vocabulary construction and TF-IDF fitting were performed within each training fold only (26).

#### Causal language detection model

We implemented and compared four supervised classifiers representing complementary modeling assumptions and complexity levels: L1- and L2-regularized logistic regression as linear, interpretable baselines (16-18); Random Forest as a non-linear ensemble method (18, 19); and Extreme Gradient Boosting (XGBoost) as a high-capacity boosting approach widely used in text classification tasks (27). All models were trained on 75% of the labeled dataset (n = 356) and evaluated on the remaining 25%, with stratified sampling to preserve class distribution (28).

#### Model specifications

Logistic regression models (L1 and L2 penalties) incorporated weights set inversely proportional to class frequencies to account for class imbalance (29). Random Forest classifiers were trained with 500 trees, mtry = √p (where p is the number of predictors), and balanced class weights (30). XGBoost was implemented with 200 boosting rounds, maximum tree depth of 4, subsample ratio of 0.8, and column subsample ratio of 0.8 per tree (27). No additional hyperparameter tuning was performed. The chosen hyperparameter settings were default settings implemented in the respective R packages outlined in Software and code availability subsection below.

#### Evaluation

Model performance was assessed using stratified 5-fold cross-validation. Evaluation metrics included accuracy, sensitivity, specificity, F1 score, positive predictive value, and prevalence detection accuracy (PDA) defined as absolute difference between the predicted prevalence and the true prevalence. To assess robustness to class imbalance, we conducted resampling-based simulations in which both the training sample size and the underlying prevalence of causal sentences were systematically varied. For each combination of prevalence (0, 0.25, 0.50, 0.75, and 1.0) and training size (75–375 sentences), labeled sentences were repeatedly resampled to construct training sets with the specified class proportions. Models were trained and evaluated under each condition, and the prevalence detection accuracy was calculated as the absolute difference between predicted and true prevalence. These simulations were conducted to verify that classifier performance was not systematically driven by the underlying prevalence of causal language rather than by the learned lexical signal (31). Ninety-five percent confidence intervals (CIs) for each metric were calculated using the standard normal approximation. Based on overall accuracy, F1 score, and prevalence detection accuracy, the best-performing model was then selected for application to the full corpus of microbiome literature. These metrics were selected to jointly evaluate overall correctness, robustness to class imbalance, and fidelity of corpus-level prevalence estimates.

#### Identification of influential words

To identify words contributing most to causal versus non-causal classification, we examined coefficients from the selected model, if appicable. Training sentences were tokenized into words, excluding tokens shorter than four characters to reduce noise (32). High-weight character n-grams were mapped back to full words using partial string matching, and coefficients were aggregated across n-grams belonging to the same word (33). The resulting summed coefficients yielded a ranked list of words, where positive values indicated association with causal language and negative values indicated association with non-causal language.

### Large-scale application to the full corpus

Following model validation, the selected classifier was applied to the remaining unlabeled abstracts. To capture variation in how conclusions are phrased, predictions were made for the last three sentences of each abstract and for the title. Predicted causal strength was classified as ‘present’ if the model’s probability estimate exceeded 0.5, a standard threshold for binary classification in probabilistic models (34).

#### Temporal, topic-specific, and country-specific analyses

For temporal analyses, documents were stratified by publication month and year, and the proportion of documents containing at least one sentence labeled as causal was calculated. For topic-specific analyses, we employed Structural Topic Model (STM) to identify thematic structures in the microbiome literature, using article titles (35). Prior to modeling, each title was tokenized into unigrams and processed to generate document-term matrices (36). STMs were trained using spectral initialization method for titles with 20 topics (k = 20) (35). The number of topics was selected based on evaluation across a range of topic numbers (multiples of five) using coherence and exclusivity metrics (37). These topics were manually inspected and labeled based on the most frequent and exclusive terms. Representative terms for each topic were identified using the FREquent and EXclusive (FREX) metric, which balances term frequency within a topic and distinctiveness relative to other topics (38). FREX highlights words that are both common within a topic and disproportionately more prevalent in that topic than elsewhere, facilitating interpretation and labeling of thematic clusters. These terms were used to assign descriptive labels to topics, such as ‘Differential abundance’ (e.g., patient, abundance, composition) and ‘Pediatric dysbiosis’ (e.g., children, dysbiosis, role).

For topic-specific temporal analyses, proportions were computed separately within each STM– derived topic distributions to examine changes in causal language within thematic areas over time. Similarly, country-specific analyses were performed by stratifying documents by the country of the corresponding author; countries with fewer than 20 publications were excluded to ensure stable estimates.

### Software and code availability

All analyses were conducted using the statistical software R (version 4.2.0) (21). The following packages were used: easyPubMed for data retrieval (39); tm (40), text2vec (41), and tokenizers (42) for preprocessing and vectorization; stm for topic modeling (35); glmnet (43), xgboost (44), and ranger (45) for experiments; caret for model training and performance assessment (46), and ggplot2 for visualizations (47). The full codebase and labeled data are available on request and archived on https://github.com/albina-tskhay/causal-language-model.

## RESULTS

### Corpus overview

A total of 20,022 abstracts published between January 2015 and July 2025 met the inclusion criteria and were included in the analysis. Overall publication volume increased steadily over the study period, reflecting the rapid growth of gut microbiome research. Leading journals in terms of total number of publications included *Scientific Reports, Gut Microbes, and Nutrients* while the countries with the largest contributions were China, Japan, and Spain (Figure 1A–E).

**Figure 1.**
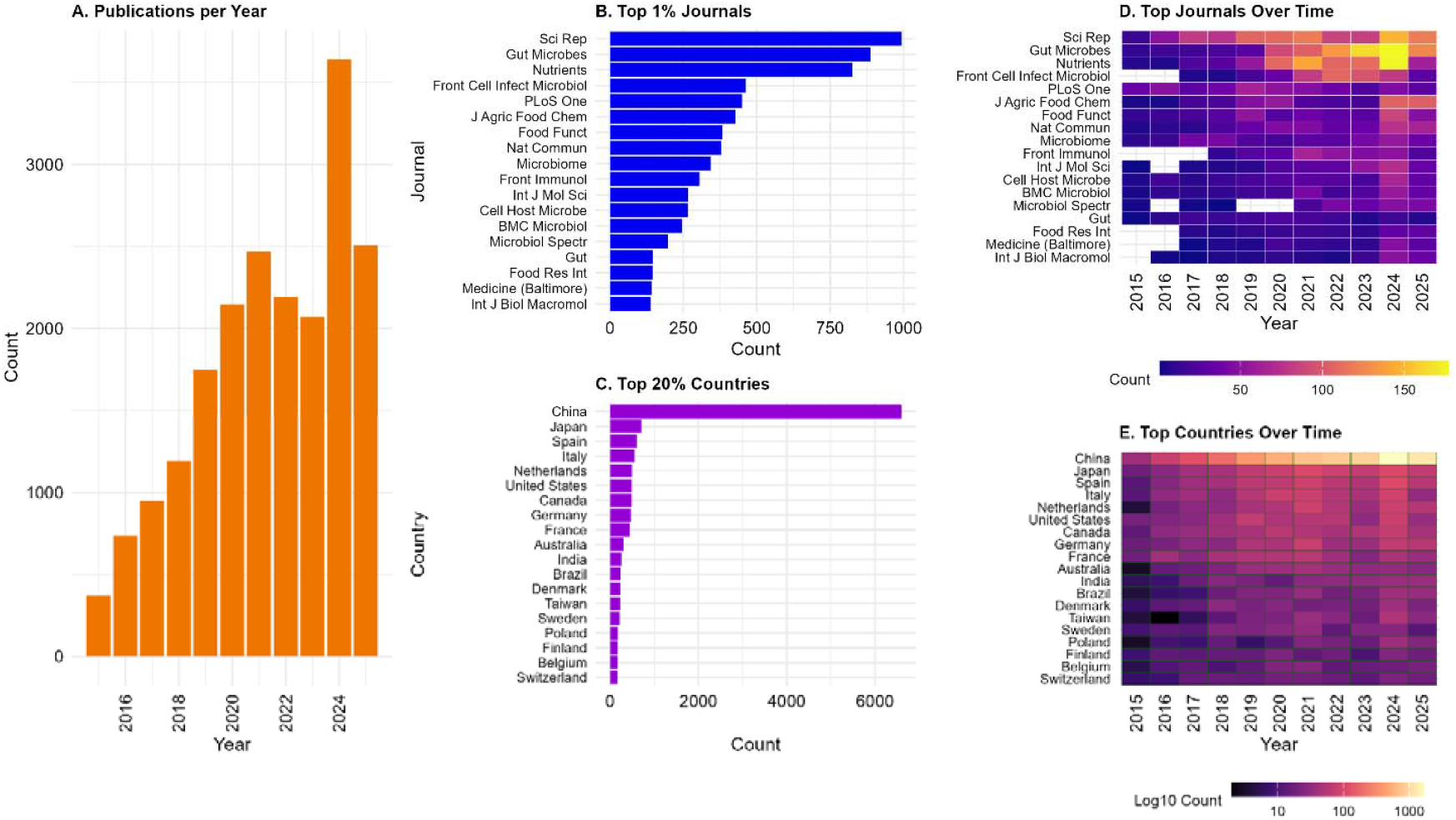
Descriptive characteristics of the microbiome literature corpus (N = 20,022). (A) Annual distribution of publications from 2007 to 2025. (B) Distribution of papers across journals, limited to the top 1% of journals by publication volume. (C) Distribution by first author’s country affiliation, restricted to the top 20% of countries by total output. (D) Temporal trends in publication output within the top 1% of journals. (E) Temporal trends in publication output within the top 20% of contributing countries.

### Model performance evaluation

With 475 sentences manually labeled, the L1-regularized logistic regression model achieved the highest performance across most metrics, with an accuracy of 76.2% (95% CI: 72 – 80%), F1 score of 72% (95% CI: 66 - 78%), and prevalence detection accuracy (PDA) of 95% (95% CI: 91– 100%) achieved on test sets (Table 1). Sensitivity and specificity were 72% (95% CI: 60 - 84%) and 80% (95% CI:72 - 87%) respectively, indicating balanced performance across causal and non-causal classes.

**Table 1.** Performance comparison of causal language classification approaches.

| **Metric** | **L-1 Logistic**<br>**regression (Lasso)** | **Random forest** | **L-2 Logistic**<br>**regression (Ridge)** | **XGBoost** |
| --- | --- | --- | --- | --- |
| **Accuracy** | 0.76 (0.72; 0.80) | 0.72 (0.68; 0.76) | 0.72 (0.68; 0.76) | 0.70 (0.67; 0.73) |
| **F1** | 0.72 (0.66; 0.78) | 0.62 (0.56; 0.68) | 0.64 (0.57; 0.71) | 0.64 (0.58; 0.70) |
| **PDA** | 0.95 (0.91; 1.00) | 0.89 (0.81; 0.97) | 0.93 (0.87; 0.99) | 0.95 (0.91; 0.99) |
| **PPV** | 0.72 (0.68; 0.77) | 0.74 (0.60; 0.88) | 0.70 (0.59; 0.80) | 0.65 (0.59; 0.70) |
| **Sensitivity** | 0.72 (0.60; 0.84) | 0.54 (0.48; 0.60) | 0.59 (0.50; 0.68) | 0.64 (0.53; 0.75) |
| **Specificity** | 0.80 (0.72; 0.87) | 0.86 (0.77; 0.94) | 0.81 (0.71; 0.90) | 0.74 (0.68; 0.80) |

Evaluation of four machine learning classifiers Metrics include accuracy, F1 score, sensitivity, specificity, positive predictive value (PPV), and prevalence detection accuracy (PDA), assessing the ability of each method to classify and estimate the prevalence of causal claims in scientific text.

Random Forest and L2-regularized logistic regression showed slightly lower overall performance. Both achieved an accuracy of 72% (95% CI: 68–76%), with F1 scores of 62% (95% CI: 56–68%) for Random Forest and 64% (95% CI: 57–71%) for L2-regularized logistic regression. Prevalence detection accuracy remained high for both models (89% [95% CI: 81–97%] and 93% [95% CI: 87–99%], respectively), indicating reasonable recovery of corpus-level prevalence despite weaker sentence-level classification. Random Forest exhibited higher specificity (86% [95% CI: 77–94%]) but substantially lower sensitivity (54% [95% CI: 48–60%]), suggesting a tendency to under-identify causal sentences. L2-regularized logistic regression demonstrated more balanced sensitivity (59% [95% CI: 50–68%]) and specificity (81% [95% CI: 71–90%]), though still inferior to the L1-regularized model across most metrics.

XGBoost achieved the lowest overall accuracy at 70% (95% CI: 67–73%) and an F1 score of 64% (95% CI: 58–70%). While its prevalence detection accuracy was comparable to that of the L1-regularized model (95% [95% CI: 91–99%]), this was driven by moderate sensitivity (64% [95% CI: 53–75%]) and lower specificity (74% [95% CI: 68–80%]), indicating less stable discrimination between causal and non-causal sentences.

Simulations varying both true causal prevalence and training size confirmed that model performance was not systematically biased by underlying prevalence (Figure S2). Mean absolute prevalence error decreased from 0.16 (50 sentences) to 0.04 (250 sentences), with only marginal gains in PDA beyond 250 sentences but narrower confidence intervals, demonstrating that the labeled dataset was sufficient for accurate corpus-level predictions. Based on these results, the L1-regularized logistic regression model was selected for application to the full corpus.

### Influential words driving classification

The L1-regularized logistic regression model was highly sparse, retaining non-zero coefficients for only 1.6% of candidate TF-IDF features (242 of 15,270). Inspection of model coefficients revealed a clear distinction between causal and non-causal language. Terms with the largest positive coefficients contributing to causal classification included suggest, increase, effect, change, and treatment alongside exposure-related terms (*antibiotic, probiotic, breastfed*) and outcome-oriented terms (*beneficial, decrease, activation, enhance*). In contrast, the strongest negative coefficients contributing to non-causal classification included *associate/association, correlate/correlation, identify, and reveal*, which typically describe observed relationships without implying directionality or intervention (Figure 2A).

**Figure 2.**
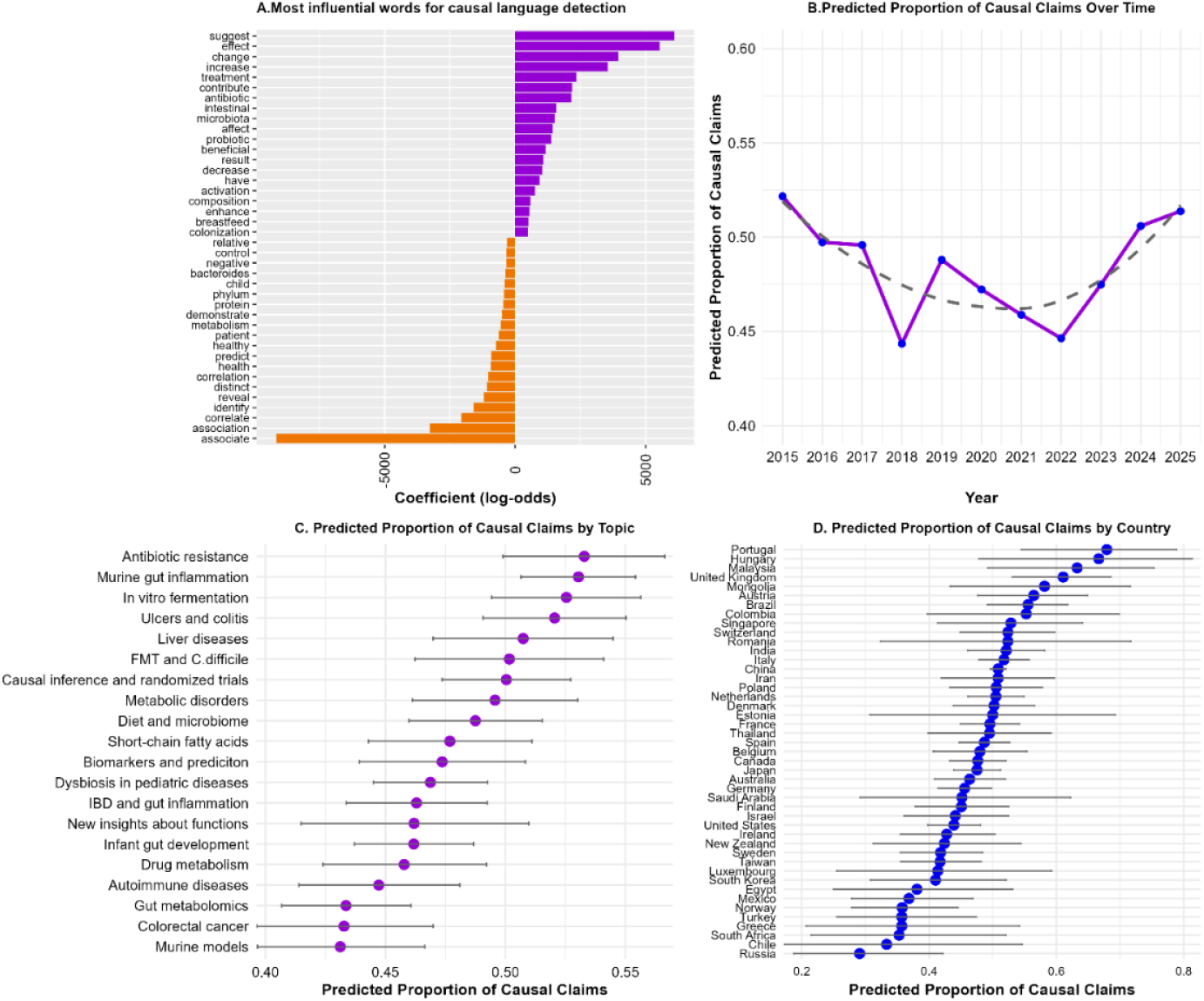
Patterns and Linguistic Signatures of Causal Claims in Microbiome Research. (**A**) Words most strongly associated with causal (positive) and non-causal (negative) language in abstracts. Importance scores are based on aggregated coefficients from an L1-regularized logistic regression on character n-grams. Only top words by absolute coefficient are shown. (**B**) Temporal trends in causal claims in microbiome abstracts. Annual proportions smoothed with LOESS show a decline from 2015–2017, followed by a gradual increase to 2025. (**C**) Proportion of causal claims across thematic subtopics with 95% confidence intervals, weighted by topic assignment probabilities. (**D**) Proportion of causal claims across countries with ≥20 publications, with 95% confidence intervals.

### Causal language prevalence in the full corpus

Application of the trained classifier to the 20,022 abstracts revealed that the prevalence of causal language fluctuated over time rather than increasing monotonically (Figure 2B). In 2015, 52.2% of abstracts contained at least one sentence predicted as causal. This proportion declined to 44.4% by 2018, stabilized between 45–47% from 2019–2022, and subsequently rose above 51% by 2025, indicating moderate variability in reporting practices despite steady growth in publication volume.

### Topic-specific patterns

Structural Topic Modeling identified 20 thematic clusters. The most prevalent topics were *Dysbiosis in pediatric diseases (10*.*3%), Gut metabolomics (7*.*3%), and Murine gut inflammation* (6.6%) (Table 2). Temporal analysis illustrated evolving thematic trends within microbiome research from 2015 to 2025 (Figure S1). Title-based analysis reflected rising interest in *Dysbiosis in pediatric diseases, Gut metabolomics, and Murine gut inflammation*, alongside decreasing emphasis on *Fecal Microbiota Transplantation (FMT) and C. difficile and Liver diseases*.

**Table 2.** Overview of Topics Identified by Structural Topic Modeling.

|  | **Topic** | **Proportion** | **FREX terms** |
| --- | --- | --- | --- |
| **1** | **Dysbiosis in pediatric diseases** | **10.3%** | **microbiota, gut, children, dysbiosis, role** |
| **2** | **Gut metabolomics** | **7.3%** | **gut, association, metabolite, profile, metabolome, serum** |
| **3** | **Murine gut inflammation** | **6.6%** | **module, mice, intestine, inflammation, regulate** |
| **4** | **Infant gut development** | **6.1%** | **infant, develop, metagenome, milk, birth** |
| **5** | **Drug metabolism** | **5.7%** | **human, metabolite, drug, impact, host** |
| **6** | **Diet and microbiome** | **5.3%** | **health, diet, obesity, adult, individual** |
| **7** | **Causal inference and randomized trials** | **5.1%** | **study, random, control, trial, mendelian** |
| **8** | **Ulcers and colitis** | **5.6%** | **cell, active, promote, colitis, ulcer** |
| **9** | **Autoimmune diseases** | **4.5%** | **multiple, arthritis, major, sclerosis, rheumatoid** |
| **10** | **Short-chain fatty acids** | **4.5%** | **intestine, alter, acid, fatty, product** |
| **11** | ***In vitro* fermentation** | **4.4%** | **colon, effect, ferment, vitro, digest** |
| **12** | **Murine models** | **4.4%** | **microbiome, model, interact, community, mouse,** |
| **13** | **Inflammatory bowel disease and gut inflammation** | **4.4%** | **disease, bowel, inflammatory, risk, patient** |
| **14** | **Biomarkers and prediction** | **4.3%** | **predict, chronic, signature, level, biomarker** |
| **15** | **Metabolic disorders** | **4.1%** | **diabetes, mellitus, type, health, starch** |
| **16** | **Antibiotic resistance** | **4.1%** | **antibiotic, resist, coli, escherichia, persist** |
| **17** | **Liver diseases** | **3.7%** | **therapeutic, drive, checkpoint, hepatocellular, urolithin** |
| **18** | **Colorectal cancer** | **3.7%** | **cancer, colorectal, diver, oral, relate** |
| **19** | **Fecal microbiota transplantation (FMT) and *C. difficile*** | **3.5%** | **transplant, infect, difficile, clostridium, recurrent** |
| **20** | **New insights about functions** | **3.4%** | **composition, function, severe, insight, new** |

Topics derived from title text. Each topic is presented with: (1) a descriptive label assigned by the authors based on semantic interpretation of top terms; (2) the proportion of documents associated with that topic, reflecting its relative prevalence in the corpus; and (3) the five most representative terms identified by the STM algorithm using FREX scoring, which balances word frequency and exclusivity.

Causal language prevalence varied across topics, ranging from 41.3% to 53.3%, with overlapping confidence intervals indicating subtle differences overall (Figure 2C). Highest prevalence was observed in Antibiotic resistance (53.3%), *Murine gut interventions* (53.0%), *In vitro fermentation studies* (52.5%), *and Ulcers and colitis* (52.0%). Intermediate prevalence was observed in *Liver diseases* (50.7%), *FMT and C. difficile* (50.2%), and *Causal inference methods* (50.1%), while lower prevalence occurred in *Colorectal cancer* (43.1%), *Murine models* (43.1%), and *Gut metabolomics* (43.4%). Causal language usage varied substantially across countries, with predicted proportions ranging from 29% to 68% (Figure 2D). Higher prevalences were observed in Portugal, Hungary, and Malaysia, intermediate levels in China, Canada, the United States, and Germany, and lower prevalences in Russia, Chilie, and South Africa.

Temporal analysis within topics revealed divergent trends. Causal language increased most in *Metabolic disorders, FMT and C. difficile, Ulcers and colitis, and Drug metabolism*, potentially reflecting evolving methodological rigor or reporting norms. Conversely, prevalence declined in *Biomarkers and prediction, Antibiotic resistance*, and *In vitro fermentation*, suggesting shifts in focus or language conventions over time (Figure 3).

**Figure 3.**
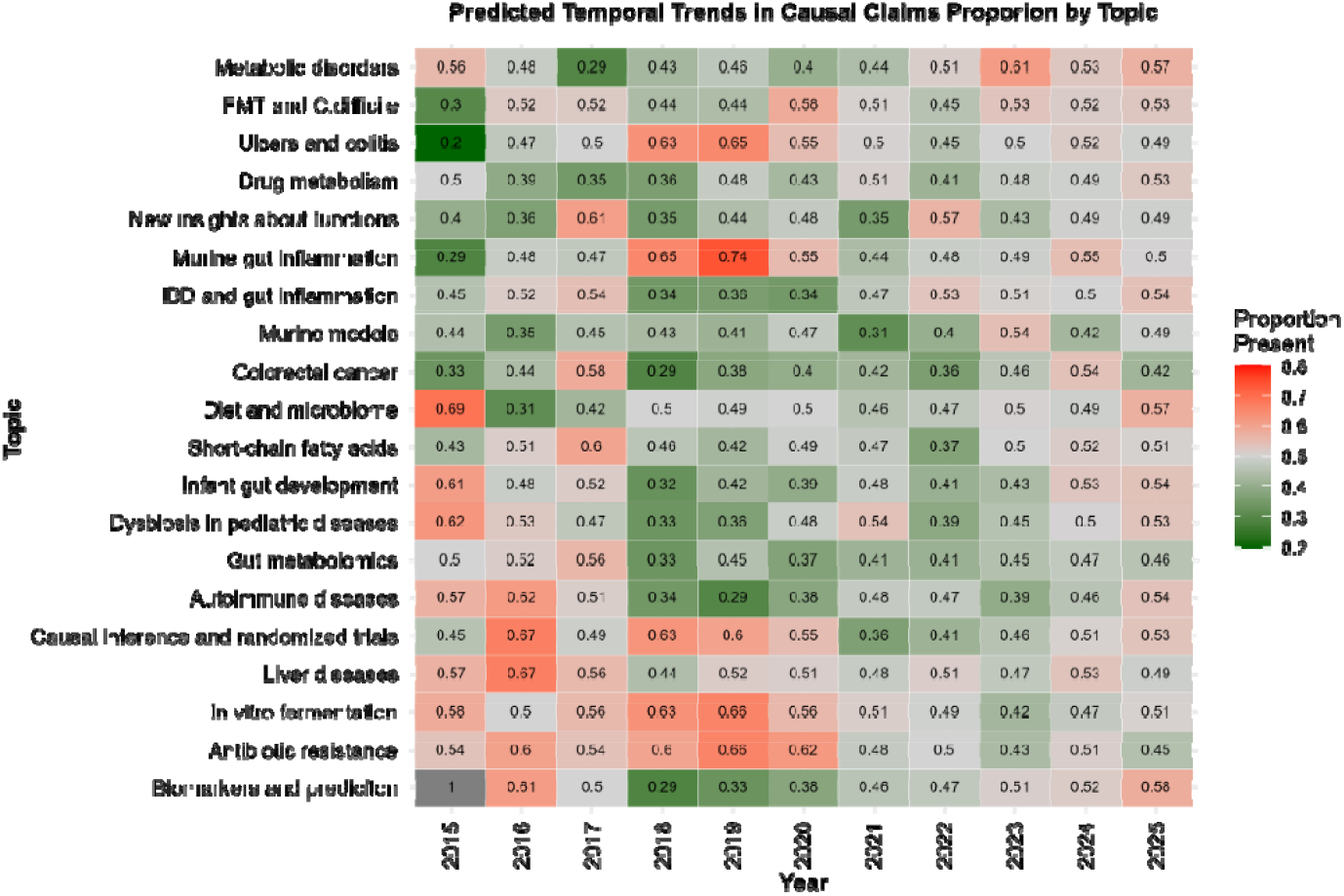
Temporal dynamics of causal claim prevalence within subtopics. This heatmap shows the temporal trends, with color intensity and numbers inside cells representing the predicted prevalence of causal claims for each topic–year combination, with red tones indicating higher prevalence and green tones indicating lower prevalence. Topics are ordered by the slope of causal language prevalence over time, with those showing increasing prevalence placed in the top rows and those with decreasing prevalence in the bottom rows.

## Discussion

This study demonstrates that a TF-IDF–based, L1-regularized logistic regression model provides satisfactory performance for detecting causal language at scale while minimizing manual annotation burden. The model likely succeeded by capturing a small set of highly influential lexical cues that help raters distinguish causal from non-causal language, including verbs and modifiers signaling change, intervention, or effect (e.g., *increase, affect, contribute*), as opposed to descriptive or associational terms (e.g., *associate, correlation, identify*). Applied to microbiome research, the approach reveals heterogeneous and evolving patterns of causal language across time, thematic subfields, and countries, with temporal dips plausibly reflecting cautious reporting and topic-specific variations.

Among the evaluated approaches, L1-regularized logistic regression consistently outperformed more complex classifiers, including tree-based ensembles such as Random Forest and XGBoost, both in overall accuracy and in estimating prevalence. Similar results have been observed in meta-research where linear models with feature selection effectively capture the “low-dimensional” nature of scientific reporting (48). This also aligns with prior work suggesting that causal language in scientific abstracts is dominated by sparse, lexically distinct constructions rather than complex interactions, making linear models with implicit feature selection particularly effective (16, 17). In comparison to human annotation, the model approaches inter-rater agreement levels documented in prior studies (4), confirming that automated detection can provide scalable insights into patterns of causal reporting. Ensemble methods, while powerful for capturing non-linear dependencies, offer limited gains in this context and may risk overfitting to idiosyncratic patterns rather than generalizable lexical cues (18, 19, 49).

The model highlights that causal meaning is primarily conveyed through a compact set of linguistic markers, consistent with earlier manual analyses. Haber et al. previously identified linguistic constructions such as “leads to,” “results in,” or “affected by” often accompanied by hedging or modality are associated with causal meaning (4). Our automated approach reproduces these findings at scale, with the dominant words overlapping those reported in manual annotation. The L1-regularized logistic regression model was highly sparse, indicating that L1-regularization ensures that only the most informative n-grams are retained, allowing the model to focus on causally salient terms while excluding language reflecting association or description. It also demonstrates that causal language detection was driven by a compact and interpretable set of lexical cues.

Applied to the gut microbiome literature, the framework provides a detailed view of how causal language is used highlighting temporal and thematic heterogeneity. The observed decline in causal reporting may reflect a period of “methodological caution” during the peak of the COVID-19 pandemic, a trend mirrored in broader biomedical literature where reporting focus shifted toward emerging observational data (7, 12). This could be explained by the fact that much of research at that time was fast-tracked and focused on immediate observational surveillance rather than long-term causal mechanisms (50).

Additionally, it was demonstrated that subfields employing experimental or interventional designs, such as *murine gut inflammation or in vitro fermentation*, exhibited higher causal language prevalence. On the one hand, this likely reflects the capacity of experimental designs to support stronger causal inference. On the other hand, when applied to clinical implementation, these patterns may suggest that the use of causal language advances faster than the adoption of rigorous causal methodologies as results derived from animal or in vitro models may not fully translate to human outcomes. For public health, this highlights potential gaps between reported claims and the underlying strength of evidence, emphasizing the need for careful interpretation when translating research findings into practice or policy (51).

In contrast, subfields dominated by observational or predictive research, including biomarker studies and colorectal cancer research, showed lower prevalence of causal language. This pattern suggests a cautious approach by authors, reflecting recognition of the inferential limitations inherent in observational data (52, 53). At the same time, it highlights opportunities for methodological improvement, such as the adoption of longitudinal designs, natural experiments, or formal causal inference methods, to strengthen the validity of claims (54, 55). Geographic patterns also revealed heterogeneity in causal language usage across countries with substantial publication output. The geographic heterogeneity observed in our study points to a “culture of reporting” that extends beyond mere data quality. Recent work by Hofstede et al. suggests that authors from countries with higher Uncertainty Avoidance Indices (UAI) or different academic training traditions may frame findings more definitively to meet perceived editorial expectations (56). This suggests that the “causal landscape” of the microbiome, and probably other biomedical research, is shaped as much by global research practices as by the underlying biological evidence indicating potential benefits of harmonized reporting standards across the global microbiome research community.

Overall, these findings have several implications for public health. When combined with full-text analysis, they could provide a framework for identifying subfields where causal claims may be over- or under-stated relative to the underlying evidence, supporting more targeted methodological scrutiny and improved reporting practices. Moreover, by linking specific linguistic cues to causal interpretation, the study offers a basis for anticipating how research findings may be interpreted in academic contexts, policy, clinical, or public health contexts. This work has several limitations. First, causal language annotation remains inherently subjective (6, 57), and abstract conclusions may omit methodological details necessary for nuanced interpretation. Second, labeled sentences were randomly split into training and testing sets at the sentence level rather than the abstract level. As a result, sentences from the same abstract could appear in both sets. While this may introduce some dependence due to shared writing style or context, each sentence was treated as an independent classification unit with distinct semantic content, because the primary goal was to assess aggregate prevalence and temporal trends rather than to optimize sentence-level prediction performance. Nonetheless, this design choice may lead to modestly optimistic performance estimates. Third, this result is based on the default hyperparameter settings, and it cannot be ruled out that the performance of other models would not be higher if the tuning was done. Fourth, while the model performs well in microbiome research, generalizability to other domains requires further validation.

## CONCLUSION

In summary, this study demonstrates that a TF-IDF–based, supervised learners can provide scalable estimates of causal language prevalence in the scientific literature. Applied to gut microbiome research, the approach reveals heterogeneous and evolving patterns of causal framing that mirror the field’s ongoing challenges in causal inference. By enabling systematic, empirical evaluation of causal language at scale, this work contributes a practical tool for monitoring scientific communication in microbiome research and beyond. Future work could extend this framework by incorporating full-text analysis, expanding labeled datasets across disciplines, or integrating causal language detection with automated assessment of study design features.

## Authors’ contributions

Albina Tskhay designed the study; labeled sentences; analyzed the data; performed the experiments, interpreted the results; and drafted the manuscript. Cristina Longo co-supervised Albina Tskhay, contributed to the design of the data extraction template, interpreted the results, and critically revised the manuscript for important intellectual content. Alibek Moldakozhayev labeled the sentences, interpreted the results, and critically revised the manuscript for important intellectual content. Nia Kang, Roxana Behruzi, Celia Greenwood, and Stan Kubow contributed to the interpretation of the results and critically revised the manuscript for important intellectual content. Tibor Schusted supervised Albina Tskhay, designed the study, interpreted the results, and critically revised the manuscript for important intellectual content.

## Human Ethics and Consent to Participate Declarations

Not applicable as no patient data were collected or analyzed in this study.

## Availability of data and materials

Code used for data processing and visualization is publicly available on GitHub at: https://github.com/albina-tskhay/causal-language-model.

## Supplementary materials

**Figure S1.**
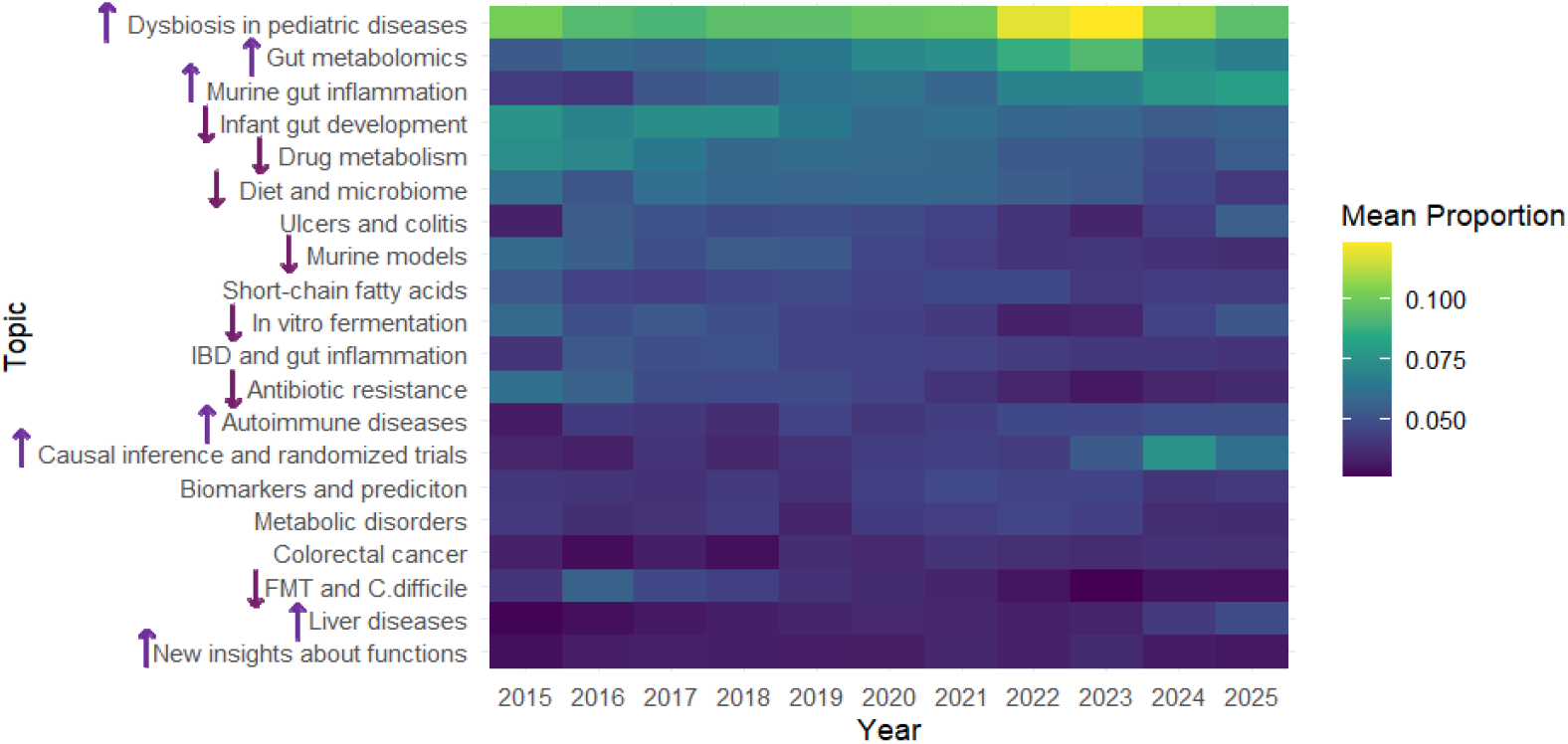
Longitudinal dynamics of topic prevalence from 2015 to 2025. Topic prevalence trends based on titles. Each line represents the average proportion of documents assigned to a given topic per year. Arrows on the left side of each panel indicate the general direction of change (increasing or decreasing) over time for easier interpretation of topic trends.

**Figure S2.**
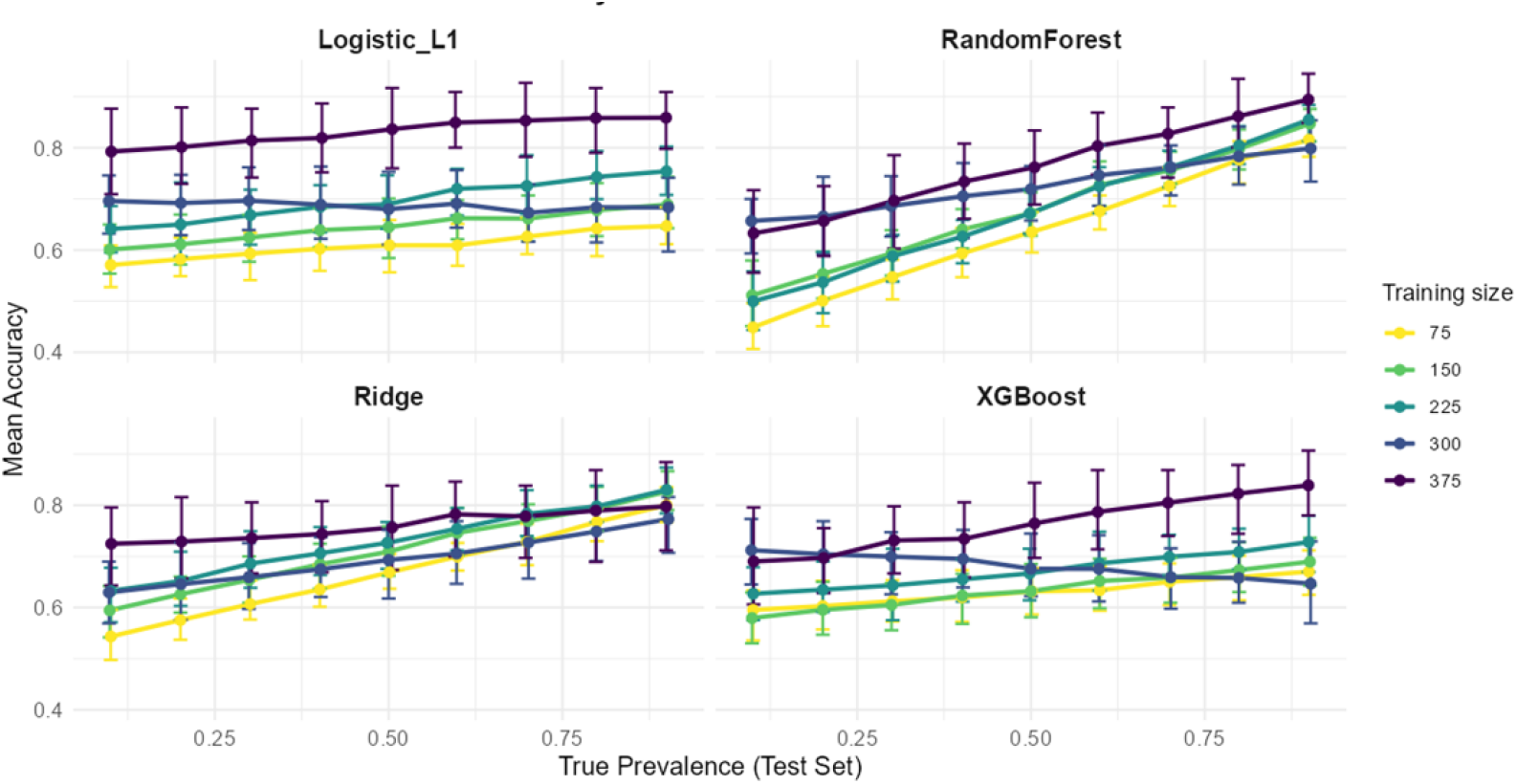
Impact of training sample size and true prevalence in the testing set on models’ accuracy. Mean accuracy in predicting causal claims with different training sample sizes and true prevalences of causal claims in testing set. As the number of labeled sentences increases, the accuracy gets higher.

